# Mechanistic and evolutionary insights into a family of aminoacyl-tRNA deacylases that protects against canavanine toxicity

**DOI:** 10.1101/2025.05.04.652006

**Authors:** Juan S. Maldonado, Stephanie Sepulveda, Sharva Karthikeyan, Karina T. Shirakawa, Ivian Merced, Alexander A. Radecki, Jordan Douglas, Wolfgang Peti, Rebecca Page, Oscar Vargas-Rodriguez

## Abstract

Aminoacyl-tRNA deacylases safeguard the accurate translation of the genetic code by hydrolyzing incorrectly synthesized aminoacyl-tRNAs. Canavanyl-tRNA deacylase (CtdA) was recently shown to protect cells against the toxicity of the non-proteinogenic amino acid canavanine, which is synthesized and accumulated by various plants. In most organisms, canavanine is ligated to tRNA^Arg^, causing translation of arginine codons with canavanine. CtdA prevents canavanine toxicity by hydrolyzing canavanyl-tRNA^Arg^. Here, we investigated the function, structure, substrate specificity, phylogenetic distribution, and evolution of CtdA. We show that CtdA is essential to prevent canavanine cytotoxicity in *Salmonella enterica*, and its heterologous expression can also rescue *Escherichia coli*. By determining the structure of CtdA, we identified its putative binding pocket and residues that modulate enzymatic activity and specificity. We also found that CtdA displays a relaxed specificity for the aminoacyl moiety substrate as it hydrolyzes arginyl-tRNA^Arg^. Finally, we showed that despite their structural homology, CtdA and the aminoacyl-tRNA hydrolytic domain of phenylalanyl-tRNA synthetase are functionally and evolutionarily divergent. Collectively, these results substantially expand our understanding of the CtdA family, providing new insights into its structure, function, and evolution. Moreover, this work highlights the diverse mechanisms unique to each organism to ensure faithful translation of the genetic code.

## Introduction

Plants are known to produce and accumulate diverse nonproteinogenic amino acids that share structural and chemical similarities with genetically encoded amino acids. These non-proteinogenic amino acids are often used as chemical defenses against herbivores, pests, or pathogens due to their toxic effects on other organisms (1-3). The toxicity of plant-sourced non-proteinogenic amino acids frequently stems from their capacity to infiltrate protein synthesis by impersonating genetically encoded amino acids (2,4-7). A prominent example is canavanine (Can), an arginine (Arg) analog produced by several legumes, including Jack bean and alfalfa (8). Ingestion or exposure to Can harms different species, including animals and bacteria (9).

The toxicity is generally caused by the inability of the translation machinery to effectively differentiate between Arg and Can, leading to the replacement of Arg residues with Can in the proteome. Can’s distinct chemical characteristics relative to Arg, including a lower isoelectric and pKa of the guanidine group (Supplementary Figure S1), destabilize protein function, resulting in cellular dysregulation (9).

The entry point of Can into proteins is via tRNA aminoacylation, where arginyl-tRNA synthetase (ArgRS), the enzyme responsible for ligating Arg to its tRNA^Arg^, fails to reject Can. Consequently, in vulnerable organisms, Can is attached to tRNA^Arg^, and the resulting Can-tRNA^Arg^ product is used by the ribosome to translate Arg codons with Can (Supplementary Figure S1) (9). To avoid self-harm, Can-producer plants have evolved ArgRSs that effectively discriminate against Can, lowering or preventing mistranslation (10). Interestingly, Can-discriminating ArgRSs have also been found in insects that feed on seeds of the Can-producer plant *Dioclea megacarpa* (11,12). In contrast, some bacteria have developed an alternative resistance mechanism involving the enzymatic degradation of Can to homoserine and hydroxyguanidine or guanidine (13,14).

Recently, a new mode of protection against Can was discovered in the bacterial species *Pseudomonas canavaninivorans and Rhizobium legumonisarum*. These organisms can use Can as a primary source of nitrogen and carbon (15). Although *P. canavaninivorans* ArgRS accepts Can, accumulation of Can-tRNA^Arg^ is prevented by a <u>c</u>anavanyl-tRNA^Arg^ <u>d</u>e<u>a</u>cylase (CtdA), which removes Can from the tRNA (Supplementary Figure S1) (16). The discovery of CtdA, a single-domain enzyme, revealed the diverse mechanisms that enable adaptation to Can-rich environments. However, despite the significance of the CtdA discovery, existing knowledge of this enzyme family is limited.

In this study, we investigated the evolution, phylogenetic distribution, structure, substrate specificity, and function of the CtdA family. Using diverse bioinformatic approaches, we define the phylogenetic distribution and evolution of CtdA and its functional relationship with the editing domain of phenylalanyl-tRNA synthetase (PheRS). We found that CtdA is widely present in bacterial genomes and encoded in some archaea and lower eukaryotic species, making it one of the few examples of free-standing aa-tRNA deacylase families found in all three domains of life. Phylogenetic studies uncovered a complex evolutionary history of CtdA, likely emerging in bacteria and later sporadically transferring to archaea and eukaryotes. We also demonstrated that the bacterial pathogen *Salmonella enterica*, one of the leading causes of food poisoning globally, encodes a functional CtdA that protects it against Can toxicity. Notably, biochemical assays show that CtdA also hydrolyzes arginyl-tRNA^Arg^, unveiling its lack of discrimination between Arg and Can. However, a mechanism involving ArgRS prevents arginyl-tRNA^Arg^ hydrolysis. Moreover, a crystal structure of CtdA revealed the putative substrate-binding pocket and helped to identify key residues that modulate its amino acid specificity and activity.

Together, these results significantly expand our understanding of the CtdA family, providing fundamental molecular, biochemical, evolutionary, and functional insights.

## Material and methods

### Cloning

The *stm2012* gene encoding CtdA (WT or mutants) from *Salmonella enterica* serovar Typhimurium 14028s was synthesized by Twist Biosciences. The *ctdA* gene was PCR-amplified with the primer pair 632.F/R and the pEVOL with 633.F/R. For cloning into pRSF, the plasmid was amplified with 596.F/R and *ctdA* with 595.F/R. For cloning into pRP1b, the plasmid was amplified with 646.F/R and *ctdA* with 645.F/R. PrimeMax polymerase (Takara) was used for PCR reactions. The DNA fragments were cleaned up and ligated using InFusion (Takara). *E. coli* Stellar chemically competent cells (Takara) were used to screen plasmid clones. Sanger DNA sequencing (Quintara Biosciences) was used to confirm clones. Oligonucleotide primers were purchased from Integrated DNA Technologies. Primers and *stm2012* gene sequences are shown in Supplementary Table S1.

### Bioinformatics analysis

Two independent HMMER searches (17) with the “Reference Proteomes” and default search options were performed using *P. canavaninivorans* or *C. perfrigens* CtdA sequences as queries. The “significant query matches” for each search were combined. Redundant results were removed together with sequence hits for full-length PheRS b-subunit to avoid misrepresentation of potential genomic CtdA repetition. Additionally, only one representative strain for each species was kept.

To build a phylogeny, the HMMER search results were curated to include samples with only sequences consisting of 180-260 residues, filtering out PheRS sequences and other possible false sequences. The resulting sequences were aligned with MAFFT v7.453 (18). The sample consisted of a taxonomically representative selection of CtdA sequences obtained by randomly selecting up to ten genera per class and one species from each selected genus. These representative sequences were further supplemented with CtdA sequences from six species of interest: *R. leguminosarum, P. canavaninivorans, Clostridium perfringens, S. enterica, Pseudomonas aeruginosa*, and *Enterococcus faecium*. This sample regime resulted in a dataset containing 226 putative CtdA sequences from across 12 bacterial, 4 eukaryotic, and 3 archaeal phyla. As an outgroup, the sample was supplemented with 19 PheRS B3/B4 domain sequences from 17 phyla across the tree of life (bacteria, archaea, and eukarya), as annotated by AARS Online (19). These sequences are specified in supplementary materials.

The phylogeny of the dataset was estimated using BEAST 2.7.6 (20), under an OBAMA site model (21), a gamma spike clock model (22), and a birth-death tree prior (23). The OBAMA method compares competing substitution models using model averaging, favoring the WAG model (24) with four gamma categories (100% posterior support) in this case. The gamma spike model is a relaxed clock model that accounts for saltative branching (i.e., an accelerated rate of evolution at the time of branching). Saltative branching was strongly favored (100% posterior support), with 15% of all evolutionary changes estimated to abruptly occur at the time of bifurcation, and the remainder occurring in a clock-like fashion along the lineages. Markov chain Monte Carlo (MCMC) convergence was diagnosed using the automated stopping MCMC package (ASM) (25), and the posterior distribution of trees was summarized using the CCD0 method (26). The BEAST 2 XML file – which contains our datasets, model specifications, and prior distributions – is available as supplementary information.

### S. enterica growth assays

The *S. enterica* serovar Typhimurium 14028s *ΔctdA* (*stm2012*) strain (27) was obtained from BEI Resources (catalog number NR-29419). *S. enterica* 14028s WT was a gift from Dr. Michael McClelland and Weiping Chu (University of California, Irvine). LB-agar plates were streaked with frozen glycerol stocks of WT and *ΔctdA* cells and incubated overnight at 37 ºC. Single colonies were used to inoculate LB media. Cells were grown overnight at 37 ºC with constant shaking.

The overnight cultures were washed twice by centrifuging for 1.5 min at 6,000 rpm, removing the supernatant, and resuspending the cells with N-minimal media (5 mM KCl, 7.5 mM (NH_4_)_2_SO_4_, 0.5 mM K_2_SO_4_, 1 mM KH_2_PO_4_, 100 mM Tris-HCl, 10mM MgCl_2_, 0.5% glycerol, and 0.1% Casamino Acids at pH 7.5). 1 μL of washed cells was used to inoculate 99 μL of fresh N-minimal media with or without L-Can (8 mg/mL). Assays with other Arg analogs, homo-Arg, homo-citrulline, and ornithine, were performed with 10 mg/mL. Growth was monitored by measuring the absorbance at 600 nm in a 96-well clear plate at 37 ºC with continuous shaking using a BioTek synergy H1 microplate reader. The mean ± SEM of four biological replicates is displayed. L-Can was purchased from Sigma.

### *E. coli* growth assays with CtdA

Single colonies of *E. coli* MG1655 or BW25113 cells harboring a pEVOL plasmid with or without the *stm2012* (CtdA) gene were used to inoculate LB media with 35 μg/mL chloramphenicol (Cm). Cells were grown overnight at 37 ºC with constant shaking. The overnight cultures were washed twice by centrifuging for 1 min at 5,000 x g, removing the supernatant, and resuspending the cells with M9 minimal media (1x M9 salts, 2 mM MgSO_4_, 0.1mM CaCl_2_, and 1% glucose). 1 μL of washed cells was used to inoculate 99 μL of fresh minimal media containing 0.1% arabinose (for induction of CtdA) and with the indicated concentration of L-canavanine. Absorbance at 600 nm (OD_600_, growth) was monitored in a 96-well clear plate at 37 ºC with continuous shaking using a BioTek synergy H1 microplate reader. The mean ± SEM of four biological replicates is displayed.

### mTyr toxicity assays

An editing defective *E. coli* MG1655 encoding a G318W genomic mutation in *pheT* (G318W) (28) was provided by Drs. Michael Ibba and Lorenzo Leiva (Chapman University). Frozen stocks of *pheT* (G318W) and MG1655 cells were used to streak LB-agar plates. Single colonies were used to inoculate LB media. Cultures were grown overnight at 37 ºC with constant shaking. The overnight cultures were washed twice by centrifuging for 1 min at 5,000 rpm, removing the supernatant, and resuspending the cells with M9 minimal media supplemented with 1% glycerol. 1 μL of washed cells was used to inoculate 99 μL of fresh M9 minimal media with or without L-mTyr (0.5 mM). Growth was monitored by measuring the absorbance at 600 nm in a 96-well clear plate at 37 ºC with continuous shaking using a BioTek synergy H1 microplate reader. The mean ± SEM of four biological replicates is displayed.

### Salmonella CtdA purification

The pRP1b plasmid harboring the CtdA gene was transformed into *E. coli* BL21(DE3) cells. A colony was used to inoculate 20-mL LB media containing ampicillin (100 μg/mL), and cells were grown overnight at 37 ºC. The overnight seed culture was used to inoculate 1-L LB media containing ampicillin. Cells were grown at 37 ºC to an OD_600_ of 0.6. CtdA overexpression was induced with 1mM isopropyl β-D-1-thiogalactopyranoside (IPTG) for 5 h at 37 ºC. Cells were harvested by centrifugation at 4 ºC and 6000 x g for 20 min. Cells were resuspended with buffer containing 50 mM Tris-Cl (pH 7.4), 150 mM NaCl, 2 mM dithiothreitol (DTT), and Complete EDTA-free protease inhibitor tablets (Roche). Lysozyme (0.75 mg/ml) was added, and the resuspended cells were incubated for 30 min on ice. The solution was sonicated for 8 min (15 seconds on and 45 seconds off cycles). The lysed cells were centrifuged for 45 min at 4 ºC and 18,000 x g. The lysate was filtered and passed through TALON Ni-NTA resin (Takara), which was pre-washed with lysis buffer. CtdA was eluted with an imidazole gradient and stored in a buffer containing 50 mM Tris-Cl (pH 7.4), 150 mM NaCl, and 40% glycerol at −20 ºC. Protein concentration was determined using the Bradford assay.

### CtdA expression and purification for crystallography

pRP1b-CtdA was transformed into BL21 (DE3) cells (Agilent) using a standard heat shock transformation protocol. Cells were grown in LB Broth with selective antibiotic (kanamycin) at 37 ºC to an OD_600_ of 0.6. The cultures were cooled down to 18 ºC before the protein expression was induced by the addition of 1 mM IPTG. The cells were harvested 16 h later by centrifugation at 6,000 x g for 10 min. The cell pellets were stored at −80 ºC until purification. Cell pellets were resuspended in ice-cold lysis buffer (50 mM Tris pH 8.0, 500 mM NaCl, 5 mM imidazole, 0.1% Triton X-100, and an EDTA-free protease inhibitor tablet [ThermoFisher Scientific]) and lysed by using high-pressure homogenization (Avestin EmulsiFlex C3). Lysate was clarified by centrifugation (45,000 x g, 45 min, and 4 ºC), and the supernatant was loaded onto a Histrap column (Cytivia) preequilibrated with Buffer A (50 mM Tris pH 8.0, 500 mM NaCl, and 5 mM imidazole). The column was washed with Buffer A, and the protein was eluted with a gradient of Buffer B (50 mM Tris pH 8.0, 500 mM NaCl, and 500 mM imidazole). Fractions with CtdA were pooled and dialyzed overnight with TEV protease (in-house) in dialysis Buffer (20 mM Tris pH 8.0, 300 mM NaCl, and 0.5 mM TCEP) to cleave the His6-tag. The tag-free CtdA protein was subjected to subtraction Ni^2+^-NTA purification, concentrated, and purified using size exclusion chromatography (SEC, Superdex 75 26/60) using SEC buffer (10 mM Tris pH 8.0, 150 mM NaCl, and 0.5 mM TCEP), which showed that CtdA is a dimer. CtdA fractions were pooled and concentrated to 37.5 mg/ml.

### CtdA crystallization and structure determination

Crystals of CtdA were grown using the vapor diffusion sitting drop method in Intelli Plate 96-3 well (Art Robbins) in 0.2 M ammonium citrate dibasic, 20% PEG 3350, and CtdA at 35 mg/ml. X-ray diffraction data of all complexes were collected using a Bruker Venture D8 system (IμS Diamond source) with a Photon III M14 detector and cold stream, and the data processed using Proteome. The datasets of CtdA were phased using molecular replacement using coordinates of CtdA generated by AlphaFold3 as a search model (29,30). All structures were completed using multiple cycles of manual building and refinement using WinCoot and Phenix Refine (30,31), respectively. Data collection and refinement statistics are reported in Table 1.

**Table 1.** Data collection and refinement statistics (molecular replacement)

| **PDBID** | **CtdA**<br>**9MKL** |
| --- | --- |
| **Data collection** |  |
| Space group | P1 |
| Cell dimensions |  |
| *a*, *b*, *c* (Å) | 72.9, 87.6, 98.3 |
| *a*, *b*, *g* (°) | 63.6, 82.1, 87.4 |
| Resolution (Å) | 21.88 – 1.98 (2.01 – 1.98) |
| *R*<sub>merge</sub> | 0.185 (0.997) |
| *I* / *sI* | 7.8 (1.1) |
| CC ½ | 0.99 (0.48) |
| Completeness (%) | 97.6 (52.1) |
| Redundancy | 6.9 (2.9) |
| **Refinement** |  |
| Resolution (Å) | 21.88 – 1.98 (2.03 – 1.98) |
| *R*<sub>work</sub> / *R*<sub>free</sub> | 0.18/0.23 (0.27/0.31) |
| Mol/ASU | 8 |
| No. atoms |  |
| Protein | 13859 |
| Ligand/ion | 0 |
| Water | 1884 |
| *B*-factors |  |
| Protein | 21.9 |
| Ligand/ion | n/a |
| Water | 29.8 |
| R.m.s. deviations |  |
| Bond lengths (Å) | 0.006 |
| Bond angles (°) | 0.805 |
| Ramachandran |  |
| Outliers (%) | 0.1% |
| Allowed (%) | 2.8% |
| Favored (%) | 97.1% |
| Clashscore | 2.34 |

<sup>a</sup>Data was collected from a single crystal
\*Values in parentheses are for highest-resolution shell

### Structural modeling and analyses

Docking models were generated using the “Structure-Based Modeling Support” in ProteinsPlus (https://proteins.plus) (32). First, potential binding pockets were identified using the DoGSite3 suite. The predicted pocket consisting of S80, L84, T104, N105, V115, G116, N170, W171, R172, and Q173 was used for docking L-Can or L-Arg using the JAMDA suite with default options selected. The ten resulting models with the best scores were individually analyzed. Only the models displaying the amino acid side chain oriented into the protein binding pocket were considered. The CtdA models with Can and Arg were superimposed using the Matchmaker tool in Chimera X (33). The Matchmaker tool in ChimeraX was also used for the structural alignment of *E. coli* PheRS (PDB ID: 3PCO) (34) and CtdA using all the Cα atoms.

### ArgRS purification

*E. coli* ArgRS was purified as previously described (35). Briefly, *E. coli* cells from the ASKA collection harboring the expression plasmid for the *argS* gene (36) were grown in LB media with chloramphenicol to an OD_600_ of 0.4, followed by induction of *argS* overexpression with 0.2 mM IPTG overnight at 25 ºC. Cells were harvested by centrifugation at 4 ºC and 6000 x g. Cells were resuspended with buffer containing 50 mM phosphate buffer (pH 8), 300 mM NaCl, 10% glycerol, and protease inhibitor tablets. Lysozyme (0.75 mg/ml) was added, and the resuspended cells were incubated for 30 min on ice. The solution was sonicated for 8 min (15-sec on and 45-sec off cycles). The lysed cells were centrifuged for 60 min at 4 ºC and 16,000 x g. The lysate was filtered and passed through a Ni-NTA column, pre-washed with lysis buffer. ArgRS was eluted with an imidazole gradient and stored in a buffer containing 20 mM phosphate buffer (pH 8), 100 mM NaCl, and 40% glycerol.

### Aminoacylation assays

*E. coli* tRNA^Arg^ (*argU*), which is 96% identical to *S. enterica* tRNA^Arg^ UCU (Supplementary Figure S2), was transcribed *in vitro* and purified as previously described (35,37,38). tRNA was radiolabeled at the 3’-adenosine with ^32^P, as described before (39). Residual Arg bound during ArgRS purification was removed as previously described (16). Briefly, 20 μM ArgRS was incubated with 20 μM tRNA^Arg^ in 50 mM HEPES (pH 7.3), 4 mM ATP, 10 mM MgCl_2_, 1 mM DTT, and 0.1 mg/mL BSA for 20 min at 37 ºC. The reaction was quenched with 2 M urea and 1 M NaCl. ArgRS was repurified using TALON resin, as described above. Aminoacylation assays were performed in 50 mM HEPES (pH 7.3), 4 mM ATP, 10 mM MgCl_2_, 1 mM DTT, 0.1 mg/mL BSA, 3 μM tRNA^Arg^ with trace amount of ^32^P-labeled tRNA^Arg^, 0.5 μM ArgRS, 5 mM Can or Arg, and 2 μM CtdA. The reactions were initiated by adding ArgRS and incubated at 37 ºC. At the indicated times, 2 μL of the reaction solution was mixed with 5 μL 0.1 units/μl P1 nuclease (Sigma) in 200 mM sodium acetate, pH 4.5. The quenched time points were incubated at room temperature for 1 h. 1 μL of the quenched solution was spotted on cellulose thin-layer chromatography (TLC) plates (EMD Millipore). The plates were run in 0.1M ammonium acetate and 5% acetic acid to separate aminoacyl-A76 from A76. The TLC plates were air-dried, and the radioactive products were detected by autoradiography using a Sapphire Biomolecular Imager (Azure Biosystems) and quantified using ImageJ (40).

### Deacylation assays

Arg-tRNA^Arg^ was prepared by incubating 1 mM Arg, 8 μM tRNA^Arg^, and 5 μM ArgRS in a solution containing 50 mM HEPES (pH 7.3), 4 mM ATP, 10 mM MgCl_2_, 1 mM DTT, and 0.1 mg/mL BSA, and 0.2 units inorganic pyrophosphatase (New England Biolabs). The reaction was incubated at room temperature for 1.5 h and quenched with 0.3 % acetic acid. Arg-tRNA^Arg^ were phenol-extracted, followed by ethanol precipitation. The tRNA was resuspended in diethylpyrocarbonate-treated water and stored at −80 ºC. Deacylation assays were carried out in 50 mM HEPES (pH 7.3), 10 mM MgCl_2,_ 1 mM DTT, and 0.1 mg/mL BSA with 2 μM CtdA or 1 μM ArgRS. The reaction time course was monitored by mixing 2 μL of reaction solution with 5 μL 0.1 units/μl P1 nuclease in 200 mM sodium acetate (pH 4.5). The plates were run in 0.1M ammonium acetate and 5 % acetic acid to separate Arg-A76 from A76. The TLC plates were air-dried, and the radioactive products were detected by autoradiography using a Sapphire Biomolecular Imager (Azure Biosystems) and quantified using ImageJ (40).

## Results

### Salmonella enterica is resistance to canavanine

Our bioinformatic search identified putative CtdA genes in several human pathogens of considerable public health interest, including *S. enterica, Pseudomonas aeruginosa, Klebsiella pneumoniae*, and *Enterococcus faecium* (41) (Supplementary Table S2). To determine if *S. enterica* is insensitive to canavanine exposure, we compared its growth in media supplemented with the amino acid. Growth curves showed that the presence of canavanine minimally slowed *S. enterica* growth (Figure 1A). Next, we used an available *ctdA* knockout strain of *S. enterica* (Δ*ctdA*) (27) to directly validate the role of CtdA in Can resistance. Notably, the Δ*ctdA* cells did not grow in Can-supplemented media (Figure 1B). To corroborate that the deletion of CtdA is responsible for the sensitivity of *S. enterica* to Can, we complemented the Δ*ctdA* cells with a plasmid harboring the *ctdA* gene. As anticipated, the expression of plasmid-borne CtdA rescued Δ*ctdA* growth in the presence of Can (Figure C-D). No effect was observed for overexpression of CtdA in *S. enterica* WT cells (Supplementary Figure S3). We also tested whether other Arg analogs could have similar toxic effects as Can in *S. enterica. S. enterica* WT and Δ*ctdA* cells were cultured with the known natural metabolites homo-Arg, homo-citrulline, or ornithine, (Supplementary Figure S4A). However, only Can diminished the ability of *S. enterica* Δ*ctdA* cells to grow (Supplementary Figure S4B-G). Together, these results demonstrate that *S. enterica* encodes a functional CtdA that protects against Can toxicity.

**Figure 1.**
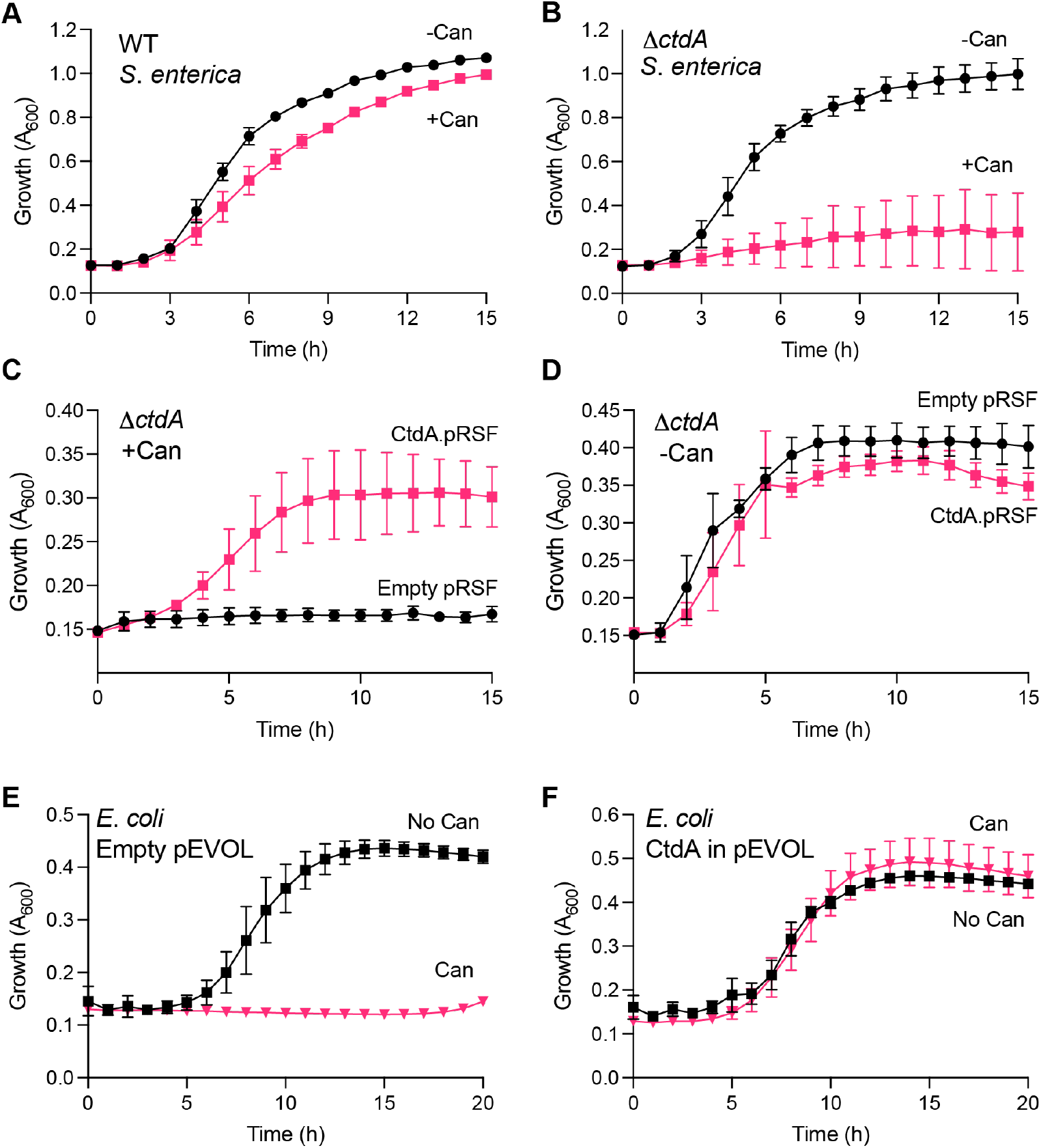
CtdA protects against canavanine (Can) toxicity. *S. enterica* WT (A) and Δ*ctdA* (A) grown in N-minimal media with or without Can. *S. enterica* WT and Δ*ctdA* cells harboring pRSF encoding or lacking CtdA were grown in N-minimal media with (C) or without (D) Can. E) and F) *E. coli* BW25113 cells carrying pEVOL with or without the *S. enterica ctdA* gene grown in M9 media without or with Can. 10 mg/mL Can was used in all experiments, and growth was monitored by measuring absorbance at 600 nm. The growth curves represent the average of at least three biological replicates with the standard deviation indicated by the error bars.

### S. enterica CtdA protects E. coli against canavanine toxicity

Unlike *S. enterica, E. coli* is sensitive to Can due to its misincorporation into proteins (42). We posed that heterologous expression of *S. enterica* CtdA could endow *E. coli* cells resistance to Can. To test this, we transformed a plasmid harboring the *S. enterica ctdA* gene into *E. coli*, and grew the resulting strain in the presence of Can. As expected, *E. coli* cells carrying a plasmid lacking the *ctdA* gene did not grow when Can was added (Figure 1E). However, cells expressing CtdA grew in the presence of Can at the same rate relative to cells grown without the amino acid (Figure 1F). These data further support the role of CtdA as a Can detoxifying factor. Importantly, the ability of *S. enterica* CtdA to support growth of *E. coli* cells in the presence of Can enabled the facile characterization of the function of CtdA in a biosafety level 1 environment.

### S. enterica CtdA displays relaxed substrate specificity

To establish *S. enterica* CtdA as a *bona fide* canavanyl-tRNA deacylase, we recombinantly purified the protein and tested its enzymatic activity *in vitro*. We first tested whether *S. enterica* CtdA prevents accumulation of canavanyl-tRNA^Arg^ using aminoacylation assays. In these assays, we used *E. coli* ArgRS and tRNA^Arg^, which are 95% and 96% identical to *S. enterica* ArgRS and tRNA^Arg^, respectively. As expected, ArgRS synthesized Can-tRNA^Arg^ efficiently.

However, Can-tRNA^Arg^ did not accumulate when CtdA was added to the reaction, while Arg-tRNA^Arg^ synthesis was unaffected (Figure 2A). The lack of depletion of Arg-tRNA^Arg^ suggests that CtdA may distinguish between Arg- and Can-tRNA, as previously proposed (16). However, the activity of CtdA towards Arg-tRNA has not been examined in the absence of ArgRS. To directly investigate the amino acid specificity of CtdA, we prepared Arg-tRNA^Arg^ *in vitro* and incubated it with CtdA. Surprisingly, the deacylation assays show that CtdA hydrolyzes Arg-tRNA^Arg^ (Figure 2B), indicating that CtdA does not effectively discriminate against Arg. The absence of Arg-tRNA^Arg^ hydrolysis in aminoacylation assays could be due to the presence of ArgRS (Figure 2A), which may protect Arg-tRNA^Arg^ from CtdA-catalyzed hydrolysis. To examine this possibility, the deacylation activity of CtdA against Arg-tRNA^Arg^ was monitored in the presence of ArgRS. The reaction was initiated by adding Arg-tRNA^Arg^ to avoid preemptive interaction with either enzyme. The results showed that in the presence of ArgRS, CtdA does not deacylate Arg-tRNA^Arg^ (Figure 2C). In contrast, adding a 5x higher concentration of tyrosyl-tRNA synthetase (TyrRS) did not prevent Arg-tRNA^Arg^ hydrolysis by CtdA (Figure 2C).

**Figure 2.**
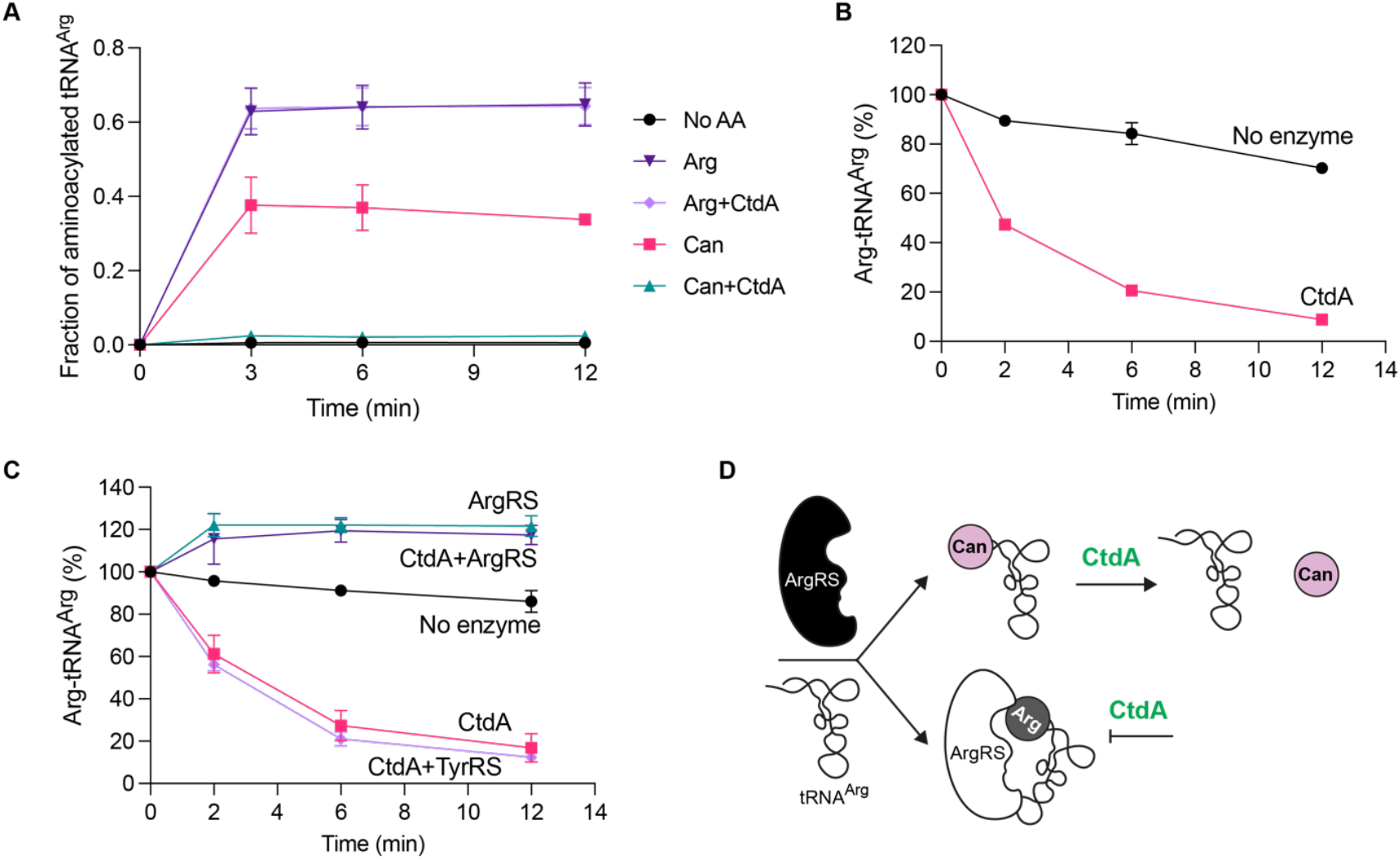
CtdA prevents Can-tRNA^Arg^ accumulation and deacylates Arg-tRNA^Arg^. A) tRNA^Arg^ aminoacylation by ArgRS (0.5 μM) with Arg or Can (5 mM) in the presence or absence of CtdA (2 μM). B) Deacylation of Arg-tRNA^Arg^ (1 μM) by CtdA (2 μM). C) Deacylation of Arg-tRNA^Arg^ (1 μM) by CtdA (2 μM) in the presence or absence of ArgRS (1 μM) or TyrRS (5 μM). “No enzyme” reactions represent Arg-tRNA^Arg^ spontaneous hydrolysis without any enzymes. The results represent the average of three independent experiments with the corresponding *standard deviation* indicated by the error bars. D) Mechanistic model of CtdA activity and specificity. CtdA deacylates Arg- and Can-tRNA^Arg^. However, CtdA is unable to hydrolyze Can-tRNA^Arg^ in the presence of ArgRS.

Interestingly, the Arg-tRNA^Arg^ concentration in the reaction slightly increased when ArgRS was added. We surmise that increased Arg-tRNA^Arg^ levels stem from the aminoacylation of free tRNA^Arg^ in the reaction and residual Arg and/or ATP and Arg-adenylate bound to the purified ArgRS. Together, these results reveal that CtdA has relaxed amino acid specificity and that ArgRS plays a crucial role in preventing Arg-tRNA^Arg^ deacylation (Figure 2D).

### Substrate recognition and catalytic mechanism of CtdA

To gain structural and mechanistic insights into CtdA’s substrate recognition and catalysis, we determined the crystal structure of *S. enterica* CtdA. CtdA adopts a B3/B4 fold that is also present in phenylalanyl-tRNA synthetase (PheRS) β-subunit (43) and is a dimer in solution (Figure 3A and Supplementary Figure S5A), with β-strand β10 residues ^224^TLIGV^228^ (monomer A) and β10 residues ^220^RFESTL^225^ (monomer B) binding in an anti-parallel fashion to extend the CtdA central β-sheet across both monomers (Supplementary Figure S5B). A DALI structural similarity search shows that CtdA exhibits the highest structural similarity to the B3/B4 domain of *E. coli* PheRS (PDBID 6p26; Dali Z-score, 15.0; root mean square deviation, 2.6 Å^2^; sequence identity, 14%; Supplementary Figure S6A) (44,45). As expected based on their structural similarities, a cavity analysis shows that the putative aminoacyl binding site is likely equivalent to that experimentally determined for Phe in the B3/B4 domain of PheRS (Supplementary Figure S6A) (46,47). Consistent with this, the base of the putative binding pocket is positively charged, as would be expected for an enzyme that binds aminoacyl-tRNA for deacylation (Supplementary Figure S6C). To test for potential substrate binding interactions, we employed docking modeling of CtdA with Can or Arg. The analyses showed that both Can and Arg are accommodated similarly in the putative binding pocket (Figures 3B and C), which may explain the relaxed enzymatic specificity of CtdA towards Arg-tRNA. The pocket is defined by several residues, including N105, G116, E118, C166, Q173, and I189. Multiple sequence alignments defined the conservation of these residues among CtdA from different species (Figure 4A Supplementary Figure S7), supporting their potential functional role in substrate recognition and catalysis.

**Figure 3.**
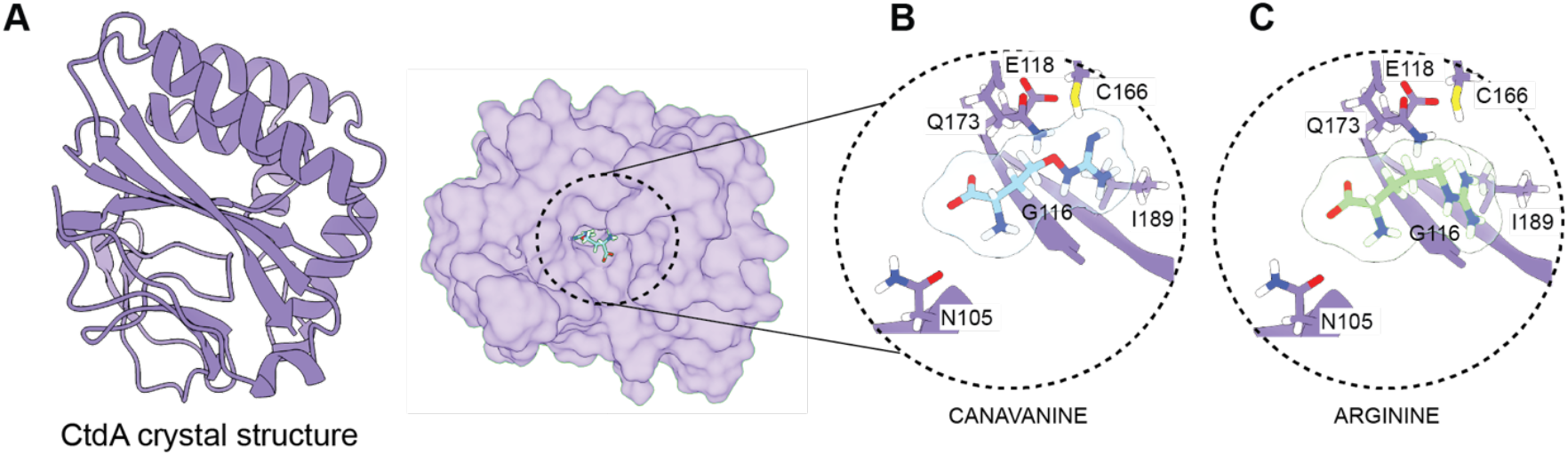
Structural models of substrate-bound CtdA. A) Crystal structure of *S. enterica* CtdA shown as a monomer. B) and C) Docking models of CtdA with Can and Arg, respectively. Several residues in the putative binding are indicated.

**Figure 4.**
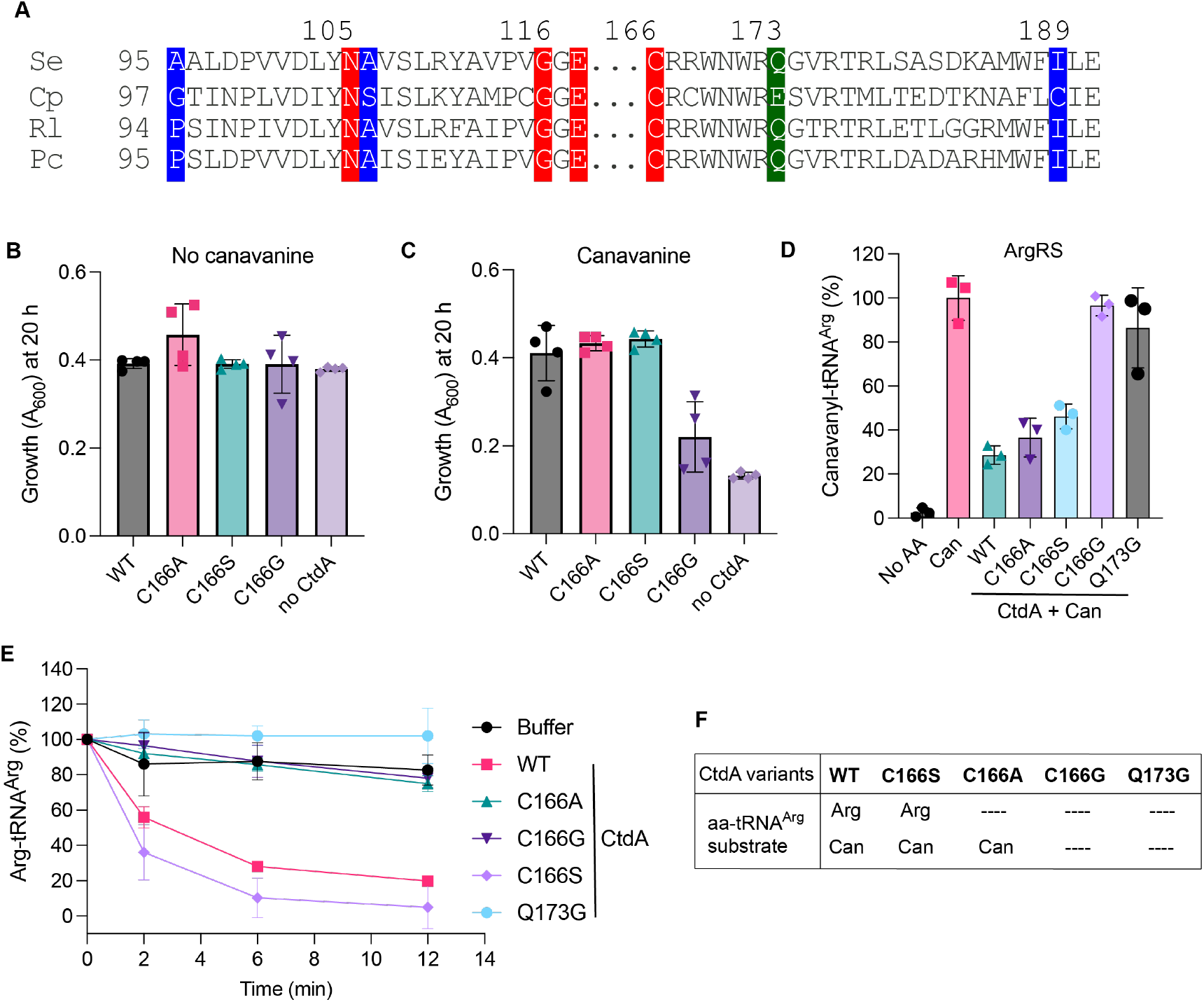
A single conserved residue modulates CtdA’s specificity. A) Multiple sequence alignments of CtdA. The highlighted residues are part of CtdA’s binding pocket. Residues highlighted in red are strictly conserved, while those highlighted in blue and green are partly conserved. The complete alignment is shown in Supplementary Figure S7. B) Growth of *E. coli* cells expressing plasmid-encoded CtdA variants after 20 h in minimal media lacking Can. C) Growth (absorbance at 600 nm) of *E. coli* cells expressing plasmid-encoded CtdA variants after 20 h in minimal media supplemented with 10 mg/mL canavanine. Each bar represents the average of four biological samples, with error bars indicating the standard deviation. Full growth curves are shown in Supplementary Figure S5). D) Aminoacylation of tRNA^Arg^ by ArgRS in the absence or the presence of Can and different CtdA variants. E) Time-course deacylation of Arg-tRNA^Arg^ by CtdA variants. The “buffer” curve indicates a reaction with no CtdA added. For D) and E), the bars represent the average of three independent trials with the standard deviation indicated by the error bars. F) Summary of the activity of CtdA variants against different Can- or Arg-tRNA^Arg^.

We were particularly interested in the role of C166 since its sulfhydryl side chain is located close to the oxa position of the Can’s side chain (Figure 3B). To investigate its role, we created three CtdA mutants (C166S, C166A, and C166G), intending to introduce distinct physicochemical and size characteristics at this position. Using our plasmid-based *E. coli* platform (Figures 1E and F), we tested the activity of the three CtdA mutants. Surprisingly, the CtdA C166S and C166A variants provided *E. coli* cells with the same growth tolerance to Can relative to CtdA WT (Figures 4B-C and Supplementary Figure S8). In contrast, cells harboring the CtdA C166G mutant were significantly sensitive to Can. To ratify their functional activities, we purified the CtdA mutants and tested their ability to prevent Can-tRNA accumulation in aminoacylation assays with ArgRS. In good agreement with our cell-based assays, CtdA C166A and C166S prevented Can-tRNA^Arg^ accumulation with similar efficiency compared to the WT enzyme, while C166G did not (Figure 4D). Next, we studied the role of C166 in the hydrolysis of Arg-tRNA.

Surprisingly, although CtdA C166S and C166G display the same activity for Arg-tRNA relative to Can-tRNA, C166A was inactive against Arg-tRNA^Arg^ (Figure 4E-F). Similarly, CtdA C166G did not deacylate Arg-tRNA (Figure 4E-F). Collectively, these results support the predicted aminoacyl binding pocket of CtdA and unveil the role of C166 in catalysis and substrate recognition. The data indicate that the residue size at position 166 is important for catalysis, while its chemistry modulates specificity.

### Functional and structural relationship of CtdA and PheRS editing domain

CtdA genes are commonly annotated as “B3/B4 domain protein” in genomic databases due to homology with the B3/B4 domain of the β-subunit of phenylalanyl-tRNA synthetase (PheRS) (Figure 5A). The PheRS B3/B4 domain is also an aa-tRNA deacylase that hydrolyzes tRNA^Phe^ acylated with Tyr or meta-Tyr (mTyr), erroneously synthesized by PheRS (28,43).

**Figure 5.**
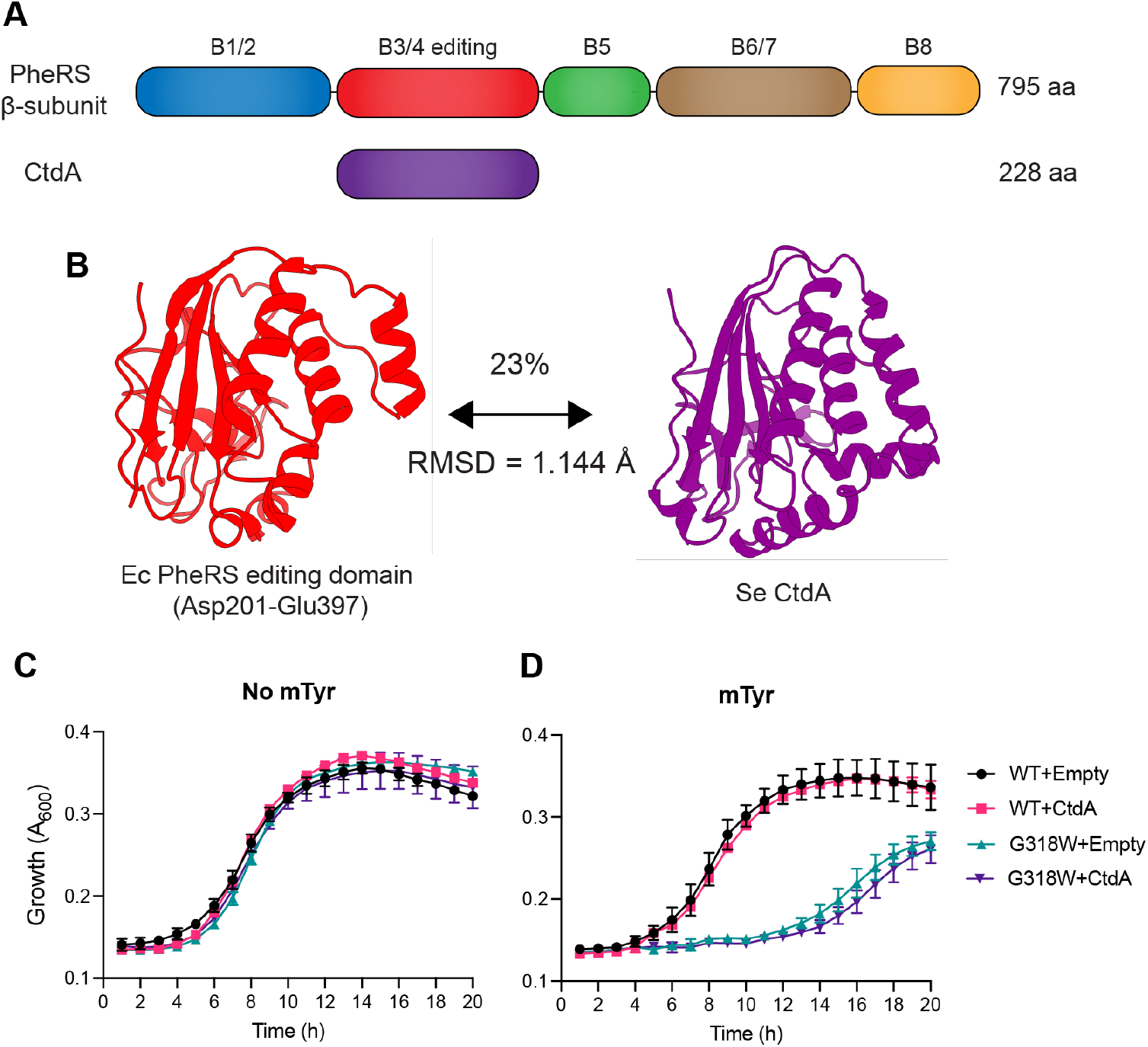
Structural and functional homology of CtdA and PheRS. A) Domain organization of PheRS β-subunit and CtdA. CtdA is a single domain homologous to the B3/B4 domain (residues Asp201-Glu397) of PheRS β-subunit, which catalyzes the hydrolysis of mTyr-tRNA^Phe^. The amino acid (aa) length of each protein is indicated. B) CtdA and the PheRS’s B3/B4 domain share low sequence identity (23%) but high tertiary structural homology (RMSD = 1.144 Å). The structures of *E. coli* (Ec) PheRS B3/B4 (PDBID 3PCO) and *S. enterica* (Se) CtdA (PDBID 9MKL) are shown as monomers. *E. coli* G318W and WT cells expressing CtdA WT from pEVOL were grown in the absence (C) and presence of mTyr (D). Growth was monitored by measuring the absorbance at 600 nm (A_600_). WT and G318W indicate WT and G318W *E. coli* cells, respectively. pEVOL lacking the *ctdA* gene (“Empty”) was used as a negative control. Each time point represents the average of three biological replicates with the standard deviation denoted by the corresponding error bar.

Consequently, the B3/B4 domain prevents translation of Phe codons with Tyr or meta-Tyr. However, beyond the genomic annotation of CtdA, little is known about the relationship between CtdA and the PheRS B3/B4 domain. To gain new insights, we performed comparative structural analyses using *S. enterica* CtdA and *E. coli* PheRS solved structures. The analyses revealed that, despite their low primary sequence homology (23%), *S. enterica* CtdA and *E. coli* PheRS B3/B4 domain share a highly conserved tertiary structure with an RMSD equal to 1.144 Å (Figure 5B). Intrigued by their homology, we asked if CtdA shares the activity of the PheRS B3/B4 domain to hydrolyze mTyr-tRNA^Phe^. To answer this question, we used an *E. coli* strain carrying the chromosomal single-residue mutation G318W in the *pheT* gene (PheRS beta-subunit). The G318W mutation is in the PheRS B3/B4 domain, and it disables its mTyr-tRNA^Phe^ deacylation activity, rendering the *E. coli* mutant strain sensitive to mTyr (Figures 5C-D) (28). To determine if CtdA could rescue *E. coli* G318W sensitivity to mTyr, we transformed *E. coli* G318W cells with a plasmid encoding CtdA and monitored growth in cultures with or without mTyr. CtdA expression did not protect the cells against mTyr toxicity (Figure 5D), demonstrating that, despite their structural homology, CtdA and the B3/B4 domain do not share functional activities.

To learn more about the structural differences that led to their divergent functional evolution, we analyzed the CtdA’s putative binding pocket and the active site of the PheRS B3/B4 domain. Our analysis found low conservation of active site residues between the two enzymes and identified stark differences at key positions (Figures 6A-B and Supplementary Figure S9). Except for a shared Asn residue, residues implicated in the B3/B4 domain substrate specificity and catalytic activity (R244, N254, Y255, H265, F267, A312, G318, and E334) (46) have different identities in CtdA. G318, one of the most consequential residues for the B3/B4 domain, corresponds to CtdA Q173. As described above, the G318 is essential for the B3/B4 domain function. Substitution of G318 with Trp decreases the binding pocket size, preventing the proper accommodation of mTyr/Tyr (28,46). To define the role of Q173 in CtdA, we generated a CtdA Q173G mutant and tested its ability to rescue *E. coli* cells from Can toxicity. The Q173G mutation renders CtdA inactive (Figure 6C), possibly due to the enlargement of its binding pocket. Another interesting difference between the two binding pockets is the identity of residue on the opposite side of G318 in the B3/B4 domain and Q173 in CtdA. A His at position 265 in the B3/B4 domain aligns with G116 of CtdA, which may compensate for creating the appropriate space for Can/Arg accommodation in the CtdA’s binding pocket. Consistent with this hypothesis, we found that replacing G116 with His or Trp inactivated CtdA (Figure 6C). We also tested the CtdA double-mutant G116H/Q173G, aiming to recreate the B3/B4 domain’s binding pocket. This CtdA mutant was also functionally defective. The only shared residue with the B3/B4 domain is CtdA’s N105. The N254 in the B3/B4 domain aids in positioning a catalytic water molecule responsible for the nucleophilic attack on the ester bond that cleaves the amino acid from tRNA (46). However, a B3/B4 domain N254A mutant retains almost complete enzymatic activity (46). N105 is predicted to also be close to the carboxylic group of the amino acid substrate, as observed in our docking models (Figure 6C). The lack of activity of an N105A mutant supports the essential role of N105 in the CtdA function (Figure 6C). Outside the binding pocket, we focused on A95. This residue structurally and sequentially aligns with R244 of the B3/B4 domain (Figure 6B). B3/B4 domain R244 is predicted to stabilize the tRNA via interaction with the phosphodiester backbone of the C75 nucleotide (46,48). A B3/B4 domain R244A variant is almost 3-fold less active than the WT enzyme (46). Whether A95 plays a similar function in CtdA is unknown. However, an A95-to-R mutation did not substantially alter CtdA activity (Figure 6C).

**Figure 6.**
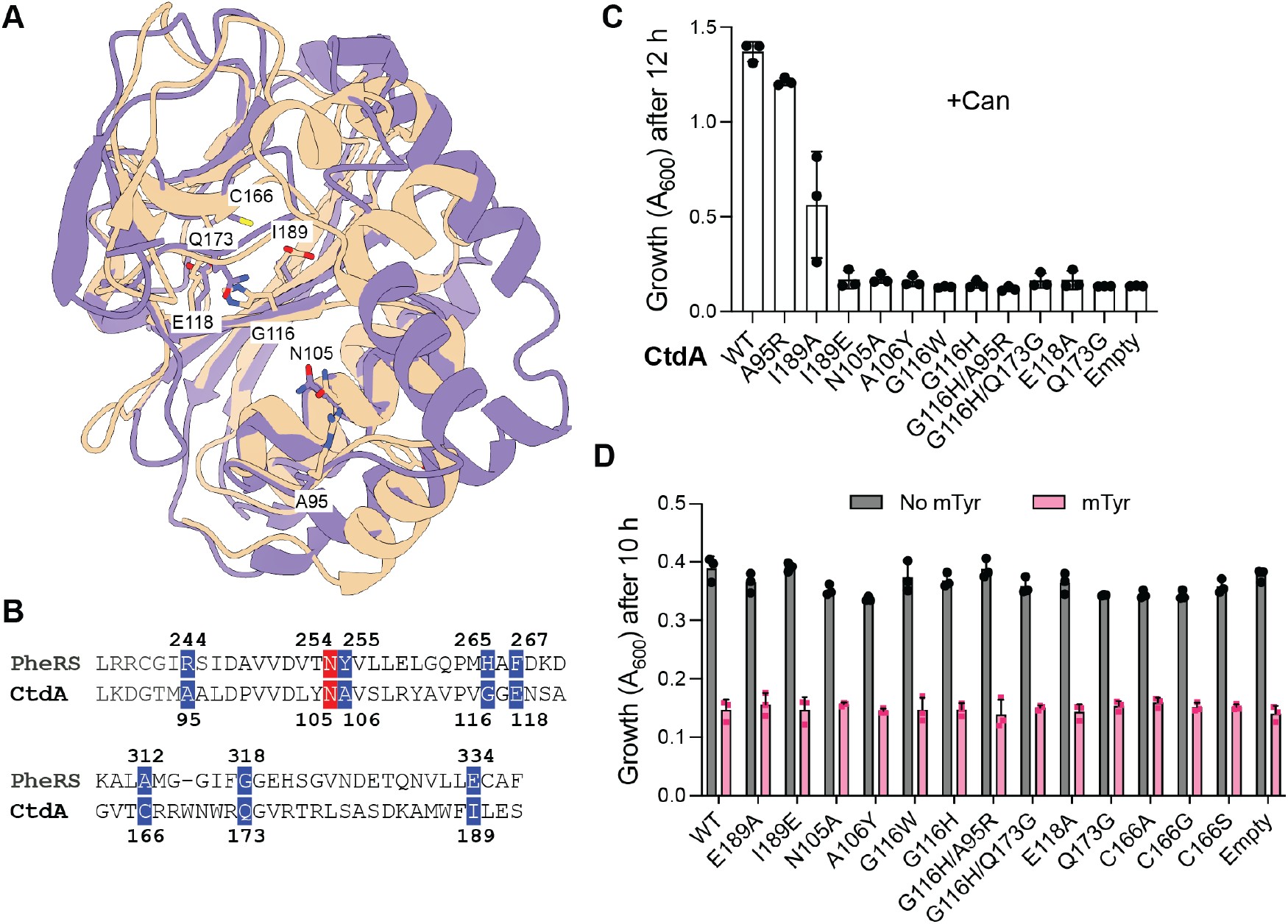
Structural and functional relationship of CtdA and PheRS editing domain. A) Structural alignment of *S. enterica* CtdA (purple) and *E. coli* PheRS editing domain (tan). Predicted active site residues for both enzymes are displayed; the residue numbers correspond to CtdA. B) Sequence alignment of *S. enterica* CtdA and *E. coli* PheRS editing domain showing the conservation of predicted active site residues (complete sequence alignment in Figure S9). C) *E. coli* BW25113 cell growth after 12 h in M9 media supplemented with Can (10 mg/mL) and pEVOL-based expression of CtdA variants induced with 0.1% arabinose. D) *E. coli* G318W cell growth after 10 h in M9 media supplemented with m-Tyr (0.5 mM) and pEVOL-based expression of CtdA variants induced with 0.1% arabinose. Bars on the graphs represent the average of three biological replicates with the standard deviation indicated by the error bars. “Empty” indicates cells carrying pEVOL without the CtdA gene.

Lastly, we explored the possibility that the mutations in CtdA could change its specificity towards mTyr, particularly because most of the mutations were made in the context of the B3/B4 domain. Also, although the mutations inactivated CtdA function against Can, the lack of activity may be caused by a change in substrate specificity rather than catalytic activity since these residues may be involved in substrate recognition but not in catalysis. We tested if any of the CtdA mutants could rescue the sensitivity of *E. coli* G318W cells to mTyr. However, none of the CtdA mutants supported *E. coli* G318W growth in mTyr-supplemented media (Figure 6D and Supplementary Figure S10), suggesting that these mutations do not change the specificity towards mTyr.

### Phylogenetic distribution of CtdA

CtdA genes were previously reported in *P. canavaninivoran, R. legumonisarum*, and *Clostridium perfringens* (16). However, the overall occurrence of CtdA in nature is unknown. To determine the phylogenetic distribution of CtdA, we performed HMMER homology searches (17) using the sequences of *P. canavaninivorans* and *C. perfrigens* CtdA as queries. These two sequences were chosen due to their low sequence similarity (∼31%), which we expected to aid in identifying diverse CtdA and CtdA-like genes more broadly across genomes. Our searches uncovered a widespread distribution of putative CtdA genes in bacteria (Supplementary Table S2). Interestingly, we also identified putative CtdA genes in archaea and eukaryotes, although they were found more sparsely. In total, 1250 organisms with reference proteomes (49) encoding CtdA were identified, corresponding to 1114 (89%) bacteria, 94 (7.5%) eukaryotes, and 42 (3%) archaea (Supplementary Table S2). In bacteria, CtdA is primarily present in Actinobacteria, Firmicutes, Bacteroidetes, and Proteobacteria (Supplementary Figure S11). In eukarya, CtdA is predominantly found in fungi belonging to the *Ascomycota* phylum, while in archaea, it is present in species from the *Euryarchaeota* and *Crenarchaeota* phyla (Supplementary Table S2 and Figure S11).

### The phylogeny and evolution of the CtdA family

We performed a phylogenetic analysis to better understand the relationship and evolution of the CtdA family. Representative PheRS B3/B4 domains from the three domains of life (50) were also included. The phylogeny of CtdA revealed two distinct CtdA clades (Figure 7A). The larger clade, which we named CtdA-1, comprises most bacterial and all the eukaryotic CtdA sequences, including the *S. enterica, P. canavaninivorans, C. perfringens*, and *R. leguminosarum* CtdA. The second smaller clade, which we named CtdA-2, consists of CtdA genes from primarily two bacterial groups (Chloroflexi and Bacillota) and all the archaeal species. The relationship of CtdA involves different instances of horizontal gene transfer. These results offer a possible path for the evolution of CtdA (Figure 7B). In this scenario, CtdA may have evolved from a B3/B4-like domain (possibly with Tyr/mTyr-tRNA hydrolytic activity), present at the time of the last universal common ancestor (LUCA). Duplication and functional diversification of the B3/B4-like domain led to the emergence of the CtdA-1 group in bacteria. It is unclear if the B3/B4 domain from which CtdA emerged was a free-standing homolog or the extant B3/B4 domain of PheRS. Members of CtdA-1 clade were later horizontally transferred to lower eukaryotes in at least two instances. A second duplication and functionalization event prompted the emergence of CtdA-2, which was later transferred to archaeal species.

**Figure 7.**
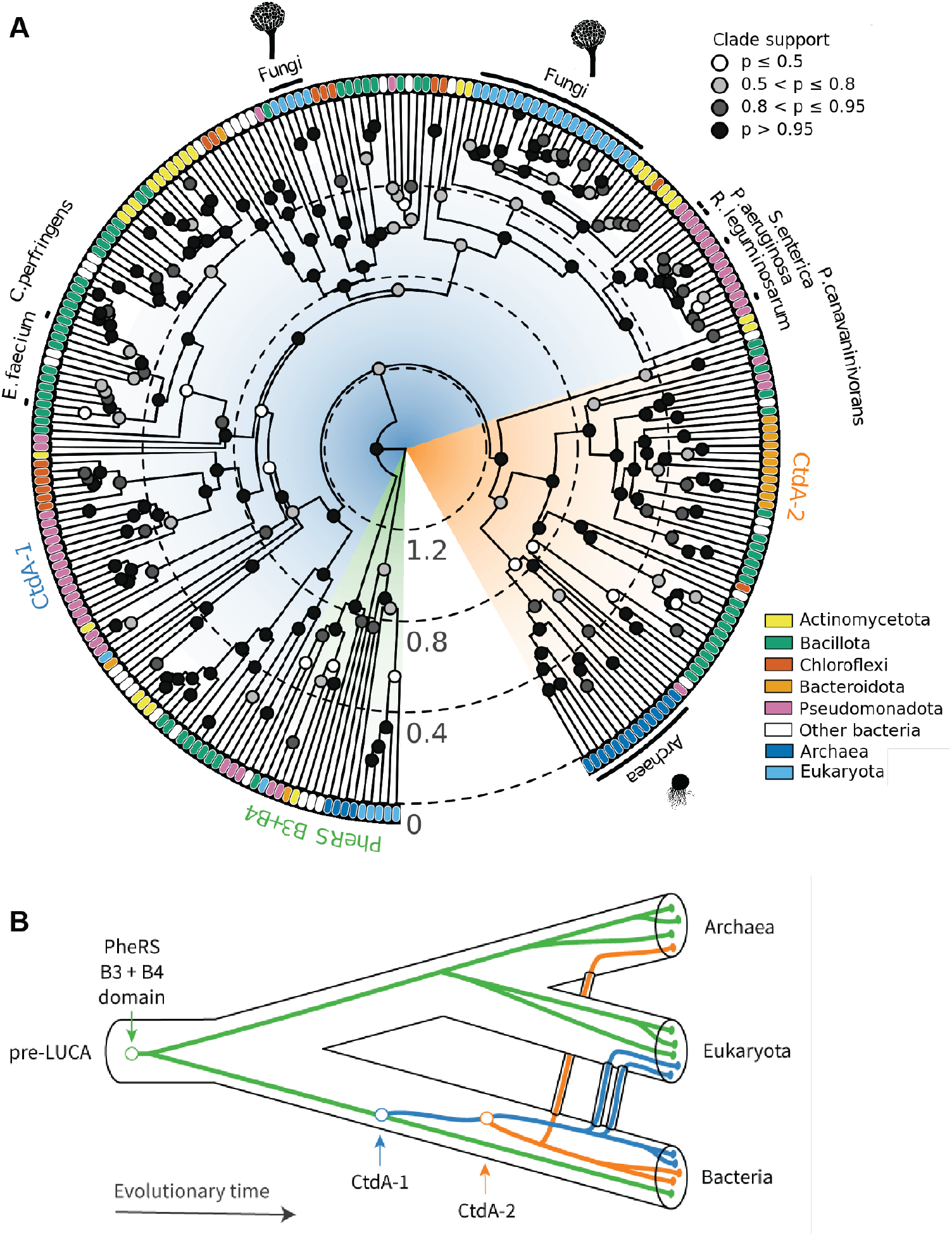
Evolution of CtdA. A) The phylogenetic tree summarizes the distribution of CtdA in organisms from all domains of life and reveals the evolution of CtdA into two distinct clades, CtdA-1 and CtdA-2. Only members of the CtdA-1 clade have been biochemically characterized. Branch lengths are in units of amino acid substitutions per site. B) An evolutionary model of CtdA. This model suggests that CtdA initially emerged from a duplication of a bacterial B3/B4-like domain, resulting in CtdA-1 proteins. A second duplication event involving bacterial CtdA-1 led to the evolution of CtdA-2. After their appearance, CtdA-1 and CtdA-2 have been horizontally transferred to eukaryotic and archaeal species, respectively. Whether the CtdA emerged from free-standing B3/B4 protein or the extant PheRS B3/B4 domain is unknown.

## Discussion

In this work, we sought to fill important knowledge gaps about the recently discovered CtdA family. Using computational approaches, we identified putative CtdA genes in over a thousand species. While predominantly found in bacteria, CtdA genes were identified in several eukaryotic and archaeal organisms. Given its phylogenetic distribution, CtdA joins a small group of standalone aa-tRNA deacylase families (ProXp-ala, AlaX, and D-aminoacyl-tRNA deacylases) with presence in the three domains of life (51). We also found that the CtdA family has evolved into two distinct groups, which we named CtdA-1 and CtdA-2. To date, only CtdAs from the CtdA-1 group have been studied. Thus, whether CtdAs belonging to the CtdA-2 group share the same functional activity or specificity is unknown.

Another poorly understood aspect of CtdA was its substrate specificity. It was initially suggested that CtdA discriminated between Can and Arg (16). However, our enzymatic studies showed that *S. enterica* CtdA also recognizes Arg-tRNA^Arg^ as a substrate. This finding questions the originally proposed naming of the CtdA family as <u>c</u>anavanyl-tRNA <u>d</u>e<u>a</u>cylase. Although it is unknown whether the deacylation of Arg-tRNA^Arg^ is biologically relevant, the potential role of CtdA’s dual specificity in other biological processes should be considered. This may help explain the presence of CtdA in organisms that, unlike *P. canavaninivorans* and *R. leguminosarum*, may not be exposed to Can in their ecological environments or use it as a metabolic source of nitrogen and carbon.

In principle, the relaxed specificity of CtdA for the aminoacyl moiety could negatively impact cellular fitness in two ways. First, unregulated hydrolysis of Arg-tRNA^Arg^ could cause undesirable ATP consumption in the aminoacylation reaction. Secondly, a decrease in Arg-tRNA^Arg^ availability could reduce protein synthesis rates (52). The relaxed substrate selectivity of CtdA is not unprecedented as other standalone aa-tRNA deacylases display relaxed specificities, resulting in the hydrolysis of correctly aminoacylated tRNAs (53). For example, human ProXp-Ala hydrolyzes Pro-tRNA^Pro^, albeit with reduced efficiency relative to its cognate substrate Ala-tRNA^Pro^ (54). Other examples of aa-tRNA deacylases with relaxed recognition include YbaK (55,56), eukaryotic AlaX (52), ATD (57), DTD, (58) and ProXp-ST1/ProXp-ST2 (59). To avoid undesirable ATP consumption and/or decreased translation rates, two major, not mutually exclusive mechanisms to prevent hydrolysis of correctly aminoacylated tRNAs are known. One mechanism involves protection of aa-tRNAs by the cognate aaRS. For example, threonyl-tRNA synthetase prevents hydrolysis of threonyl-tRNA^Thr^ by the Thr/Ser-tRNA deacylase ProXp-ST2 (59). The second mechanism involves the elongation factor (EF-Tu in bacteria or EF1A in archaea and eukaryotes), which can also protect cognate aa-tRNAs from deacylation (57,58,60). In the context of CtdA, our results support the role of ArgRS in protecting Arg-tRNA^Arg^ from CtdA-mediated deacylation. Whether EF-Tu also provides a second line of protection needs further investigation.

With the help of a new crystal structure and functional assays, we also dissected aspects of CtdA’s molecular mechanism of substrate recognition and catalysis. Docking models revealed that the putative amino acid binding pocket of CtdA can accommodate Can and Arg in a similar fashion. Notably, we found that a single residue (Cys166) can modulate the specificity of CtdA. How Cys166 modulates CtdA specificity is unclear. However, based on activities of the CtdA C166S, C166A, and C166G, we speculate that the size, rather than the chemistry, of the side chain of Cys166 is the major determining factor. Thus, enlarging CtdA’s substrate pocket by decreasing the size of residue 166 destabilizes substrate binding. Consequently, CtdA with Ser, 12 Å smaller in molecular volume than Cys (61), at position 166 retains activity and specificity. In contrast, Ala, with a 23 Å smaller molecular volume than Cys (61), at the same position weakens the binding of CtdA for Arg but not Can, while Gly disrupts the binding of both amino acids. Similar strategies of substrate recognition have been reported for the editing domain of ProRS (62) and the free-standing aa-tRNA deacylase, ProXp-Ala (54,61). Additional biochemical and biophysical evidence is needed to better understand the CtdA’s specificity and role of Cys166.

The discovery of C166 as a modulator of specificity and catalysis is also intriguing from an evolutionary standpoint. Clearly, a single residue substitution could redefine CtdA’s specificity for Can-tRNA, raising the question of why CtdA retains a relaxed specificity for the aminoacyl moiety. One possibility is that the extant CtdA is an evolutionary intermediate. Consequently, CtdA may evolve to be specific for Can by replacing C166 with Ala. An alternative scenario is that evolutionary pressure supports CtdA’s relaxed specificity, endowing it with additional biological roles beyond Can detoxification that remain unknown, including the potential activity against other amino acids. The latter case may help explain the presence of CtdA in organisms that may not be exposed to Can or do not use it as an energy source. Future studies may provide support for either hypothesis.

## Data availability

The structure of *S. enterica* CtdA was deposited in the Protein Data Bank with accession code 9MKL. All other data are contained within the manuscript.

## Supplementary data

Supplementary Data are available at NAR Online.

## Acknowledgments

We thank Drs. Michael Ibba and Lorenzo Leiva (Chapman University) for the *E. coli* PheRS editing defective strain. The *Salmonella enterica* 14028s was a kind gift from Drs. Michael McClelland and Weiping Chu (University of California, Irvine). We also thank Dr. Steven Z. Chou and Bethlehem Abebe for technical support with the purification of CtdA, as well as Dr. Meng Choy and Ms. Brooke Dreyer for initial work on CtdA crystallization.

## Funding

This work was supported by start-up funds from the Department of Molecular Biology and Biophysics at the University of Connecticut School of Medicine to O.V.-R. J.D. was supported by the Alfred P. Sloan Foundation Matter-to-Life program (grant number G-2021-16944). This work was also supported by funds from the University of Connecticut School of Medicine to R.P. and W.P.

## Conflict of interest statement

The authors declare no conflict of interest.

## Notes

### Competing Interest Statement

The authors have declared no competing interest.

